# Deepprune: Learning efficient and interpretable convolutional networks through weight pruning for predicting DNA-protein binding

**DOI:** 10.1101/729566

**Authors:** Xiao Luo, Weilai Chi, Minghua Deng

## Abstract

Convolutional neural network (CNN) based methods have outperformed conventional machine learning methods in predicting the binding preference of DNA-protein binding. Although studies in the past have shown that more convolutional kernels help to achieve better performance, visualization of the model can be obscured by the use of many kernels, resulting in overfitting and reduced interpretation because the number of motifs in true models is limited. Therefore, we aim to arrive at high performance, but with limited kernel numbers, in CNN-based models for motif inference.

We herein present Deepprune, a novel deep learning framework, which prunes the weights in the dense layer and fine-tunes iteratively. These two steps enable the training of CNN-based models with limited kernel numbers, allowing easy interpretation of the learned model. We demonstrate that Deepprune significantly improves motif inference performance for the simulated datasets. Furthermore, we show that Deepprune outperforms the baseline with limited kernel numbers when inferring DNA-binding sites from ChIP-seq data.

## BACKGROUND

Determining how proteins interact with DNA to regulate gene expression is essential for fully understanding many biological processes and disease states. Many DNA binding proteins have affinity for specific DNA binding sites. ChIP-seq combines chromatin immunoprecipitation(ChIP) with massively parallel DNA sequencing to identify DNA binding sites of DNA-associated proteins(Zhang et al., 2008). However, DNA sequences directly obtained by experiments typically contain noise and bias. Consequently, many computational methods have been developed to predict protein-DNA binding, including conventional statistical methods(Badis et al., 2009; Ghandi et al., 2016) and deep learning-based methods(Alipanahi et al., 2015; Zhou and Troyanskaya, 2015; Zeng et al., 2016). Convolutional neural networks (CNNs) have attracted attention for identifying protein-DNA binding motifs in many studies.(Zhou and Troyanskaya, 2015; Alipanahi et al., 2015). Genomic sequences are first encoded in one-hot format; then, a 1-D convolution operation with 4 channels is performed on them. For conventional machine learning methods, the sequence specificities of a protein are often characterized by position weight matrices (PWM)(Stormo, 2000). PWM has a direct connection to CNN-based model since the log-likelihood of the resulting PWM of each DNA sequence is exactly the sum of a constant and the convolution of the original kernel on the same sequence from the view of probability model(Ding et al., 2018). Zeng et al.(Zeng et al., 2016) experimented with different structures and hyperparameters and showed that the convolutional layers with more kernels could obtain better performance. They also showed that training models with gradient descent methods is sensitive to weight initialization, showing, in turn, that training could be obstructed at local optimum of loss function. However, the use of too many kernels could introduce too much noise and, thus, overfitting, leading to misinterpretation of the model. By visualizing the recovery of the underlying motifs in the models, we found that only the several best-recovered motifs, in the sense of information content, could be equated to the true motifs, demonstrating that most kernels only act during the process of training by increasing generalization ability in order to overcome the local optimum problem(Du et al., 2018). Such kernels can be termed auxiliary kernels, and these kernels produce noise and reduce performance at the end of training. Neural networks with circular filters(Blum and Kollmann, 2019) can address this problem, but performance was only found to significantly improve in the one-kernel CNN-based model. However, since some proteins likely bind multiple motifs in the DNA sequence in omics data, the one-kernel CNN-based model cannot meet the needs of motif finding. Moreover, its overall performance is lower than expected when kernel number is limited(e.g. 16). Luo et al.(Luo et al., 2019) replaced global max pooling with expectation pooling, which is shown to increase the robustness for kernel numbers. However, expectation pooling only increases model robustness; it does not limit kernel numbers.

In contrast, neural network pruning can reduce kernel numbers and by doing so, improve inferential performance without harming accuracy in the field of computer vision (Han et al., 2015a). Pruning methods can be classified into structured and unstructured. The former refers to pruning at the level of channels, or even layers, for which the original network structure is still preserved (Li et al., 2016; Changpinyo et al., 2017; Hu et al., 2016; He et al., 2017). The latter includes individual weight pruning. Han et al.(Han et al., 2015b) developed a method whereby network weights of small magnitude were pruned, and it was very successful in highly compressed neural network models (Han et al., 2015a). Unstructured pruning can ensure that models will achieve sparse weight matrices which result in compression and acceleration with dedicated hardware (Han et al., 2016).

With evidence that models with only a few kernels can fit the PWM model very well, we propose a novel model, termed Deepprune, which utilizes pruning techniques in motif inference. Several assumptions underlie the design of Deepprune. First, by its stronger representation and optimization power, we believed that starting with training a large and over-parameterized network could provide a model with high performance. Second, for the PWM model, which often characterizes sequence specificities, several kernels which are viewed as motif detectors are enough for motif inference. Third, the inclusion of too many auxiliary kernels leads to misinterpretation of the model. Fourth, auxiliary kernels may produce noise and lower performance at the end of training. If the PWM model characterizes sequence specificities and if no interaction among different motifs is considered, then Deepprune achieves better performance with fewer kernels, markedly exceeding baseline in simulated datasets. In spite of the uncertainty of the true model, Deepprune still arrives at better performance with the same kernel numbers in real datasets, which shows the superiority of our model. Our model can also find more accurate motifs by model visualization and eliminate auxiliary kernels. All coding utilized to implement Deepprune and all the figure reproductions in the paper is available at https://github.com/klovbe/Deepprune

## METHODS

### Detecting sequence motifs with CNN

We adopt the simplest model in DeepBind as our basic neural network architecture(Alipanahi et al., 2015). The sequences are represented as numerical vectors. Each of the four nuleotides is denoted as one of the four one-hot vectors [1, 0, 0, 0], [0, 1, 0, 0], [0, 0, 1, 0], and [0, 0, 0, 1]. Consequently, a sequence *X* = *X*_1_,…, *X_L_* is transformed into a 4× *L*matrix *S*. We first add a 1-D convolutional layer with rectified linear units(ReLU) activation serving as a motif scanner layer(Radford et al., 2015), followed by a global max pooling layer. Then we add a mask layer to prune the weights according to some given criterion, which will be introduced in the next section. The last layer is a fully connected layer with sigmoid activation the output of which is the probability of a sample being positive.

Formally, if the convolutional kernels are denoted by 4×*L_F_* matrices *F*^1^*, F*^2^,…, *F^d^*, in which *L_F_* is the length of the kernel, we have

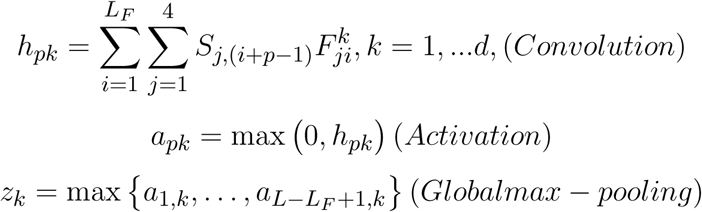

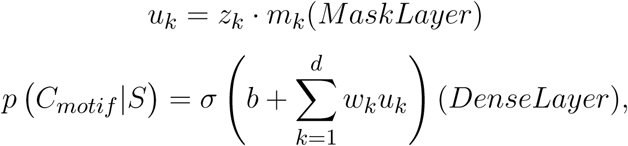

where *w_k_* and *w* are weights, *b* is bias and *σ*(*x*) denotes the sigmoid function for classification. Compared to basic neural network architectures, note that a mask layer is added because we want to mask the kernels that have little impact on the performance at the end of training. As a result, *m_k_* is set as 0 or 1, and *m_k_* = 0 means that the information of the *k*-th kernel cannot pass through this layer. Because the calculation of each kernel is independent in the convolutional layer, the pruned model can be viewed as a CNN-based model with fewer kernels. Accordingly, we can prune our network to get an efficient and interpretable architecture with limited kernels.

### Deepprune

In this work, we take iterative pruning on the weights of the dense layer in the CNN-based model and drop the learning rate of each pruning step gradually for fine-tuning. First, we utilize 2*^k^ × d* convolutional kernels in our model, i.e., the large, over-parameterized model. Half the number of kernels is pruned each time, according to a certain criterion. In other words, the number of values being 1 in the mask layer is halved each time. Since weight pruning may lead to decreased performance, we then fine-tune the pruned model to regain the lost performance. The above two steps are iterated for *k* times and then the final model is obtained. Deepprune first gives the weights in the architecture an appropriate area from the global view and adjusts the weights gradually by iterative pruning and fine-tuning. In this way, we can overcome the drawback of easily stopping at the local optima restricted by the local views in the original model with limited kernel numbers by the strong ability of representation in our model.

Three criteria are designed for Deepprune. For weight-based Deepprune, we consider the weight of scores (i.e., *w_k_*) in the dense layer. The weights with small magnitude are pruned as

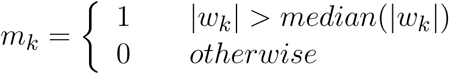

in which the median operation takes the median of *|w_k_|* corresponding to unpruned weights. However, the scale problem below is not considered in the first criterion. We know that *b* + ∑_*k*=1_^*d*^*w_k_u_k_* is the input for the sigmoid activation layer which predicts the label; that is to say *w_k_u_k_* determines the importance of the k-th kernel. However, the score of the k-th kernel can be multiplied by *m* if weights in the convolutional layer are multiplied by *m*, and then the weight corresponding to this kernel in the dense layer will shrink by training. As a result, the score *u_k_* obtained in the mask layer also counts, and the impact of the score over samples needs to be considered. For the score-based criterion, the scores with small difference between positive and negative samples are pruned.

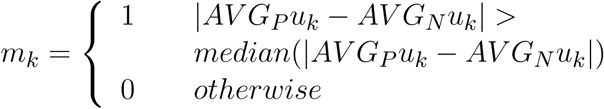

in which *AV G_P_u_k_* means the average score over positive samples, and *AV G_N_u_k_* means the average score over negative samples. For the score-and-weight-based criterion, we directly consider *w_k_u_k_*, which determines the input for the sigmoid activation layer as

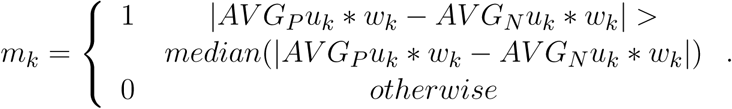

### Implementation of the models

The hyperparameters to train the simulated datasets contain the length of convolutional kernels, learning rate, times of pruning *k*, last pruned kernel number *d*, number of epochs, training batch size, learning rate decay schedule and the optimizer. First, we train the basic model with 2*^k^*×*d* kernel numbers, and we get Deepprune models with 2*^k−^*^1^×*d,…, d* kernel numbers. We also consider the strength of fine-tuning and denote the pruned model without fine-tuning from the last pruned model (twice the kernel numbers) as Deepprune-inter. To make a comparison, we match our model with baseline, which is the basic model utilizing identical kernel number trained directly without pruning.

For training, we used cross-entropy as a loss function without any weight decay(i.e., *L*_2_ regularization over the weights), and trained the model utilizing the standard backpropagation algorithm and the Adam optimizer(Kingma and Ba, 2014). The area under the ROC (AUC)(Fawcett, 2004; Davis and Goadrich, 2006) is utilized to assess prediction performance.

Our model is implemented with Keras for Python (Chollet et al., 2015).

### Datasets

#### Simulated datasets

For simulation, TRANSFAC database was utilized to evaluate the performance of Deepprune(Wingender et al., 1996). Each simulated data set includes both negative and positive samples, or sequences. Each negative sample consists of independent and identically distributed nucleotides obeying a multinomial distribution with the probability of 0.25 for each {*A, C, T, G*}. Each positive sample was built in the same manner as a negative sample except that sequences from certain motifs were inserted at some locations randomly. The sequences inserted in the positive samples for the five simulated data sets were listed below:

- **simulated dataset 1,2,3**: Each sequence was generated from either the first or the second motif; We chose motif for each positive sample randomly with equal probability.
- **simulated dataset 4**: Each sequence was generated from one of the four given motifs; other rule is the same.
- **simulated dataset 5**: Each sequence was generated from one of the eight given motifs; other rule is the same.

The number of sequences in the training dataset and test dataset is equal. We emphasized because a given protein may bind to multiple motifs in the DNA sequence, our simulation datasets were constructed reasonably.

#### Real datasets

690 ChIP-seq ENCODE datasets utilized by DeepBind were chosen to be real datasets(Alipanahi et al., 2015). Each dataset corresponds to a specific DNA-binding protein. Its positive samples are 101 bp DNA sequences confirmed to bind to a given protein experimentally while its negative samples were constructed through shuffling dinucleotides in the positive sequences. All the datasets are available at http://cnn.csail.mit.edu/.

## RESULTS

### Deepprune performs better than the baseline on the simulated data

In this section, we use the simulated data to compare Deepprune with baseline. Baseline is the simplest CNN model with no hidden layers, in other words the architecture of Deepprune, but without the mask layer, with batch size = 256, *d* = 4 and *k* = 6. All the models in this paper are pruned from the basic model with kernels [Table 1]. We chose *d* = 4 for 101 bp sequences, which can be divided into about 4 parts of 24 bp. If *d <* 4, the kernel number may be less than the number of the underlying motifs. Also, simulated dataset 5 can show how Deepprune performs when the kernel number is half the number of the underlying motifs. Several random seeds are set to evaluate the robustness of the models’ performance for the simulated datasets. Weight-based Deepprune is only considered in this section.

**Table 1.** Parameter settings for the simulated datasets

| Name | Values |
| --- | --- |
| Batch size | 256 |
| Kernel length | 24 |
| Optimizer | Adam with initial learning rate 0.01 |
| Learning rate decay schedule | drops the learning rate by 1.2 every pruning step |
| Random seed | 0,1,2,3,4,5,6,7,8,100,123,1000,1234,10000,12345,100000,123456,1000000 |

Compared to the baseline model without pruning, we found that Deepprune improved motif inference performance on first three simulated datasets from Figure 2. Specifically, as kernel number increases, the performance of baseline has a tendency to improve, which is consistent with Zeng et al.(Zeng et al., 2016) However, as kernel number decreases, the performance of Deepprune shows a converse tendency such that the mean of AUC of Deepprune shows significant improvement as the iteration continues. What’s more, variances of AUC of Deepprune are also more robust. When compared with models with the same kernel number, Deepprune shows its wonderful ability to limit kernels for accuracy and robustness, showing that Deepprune works effectively for motif inference.

**Figure 1.**
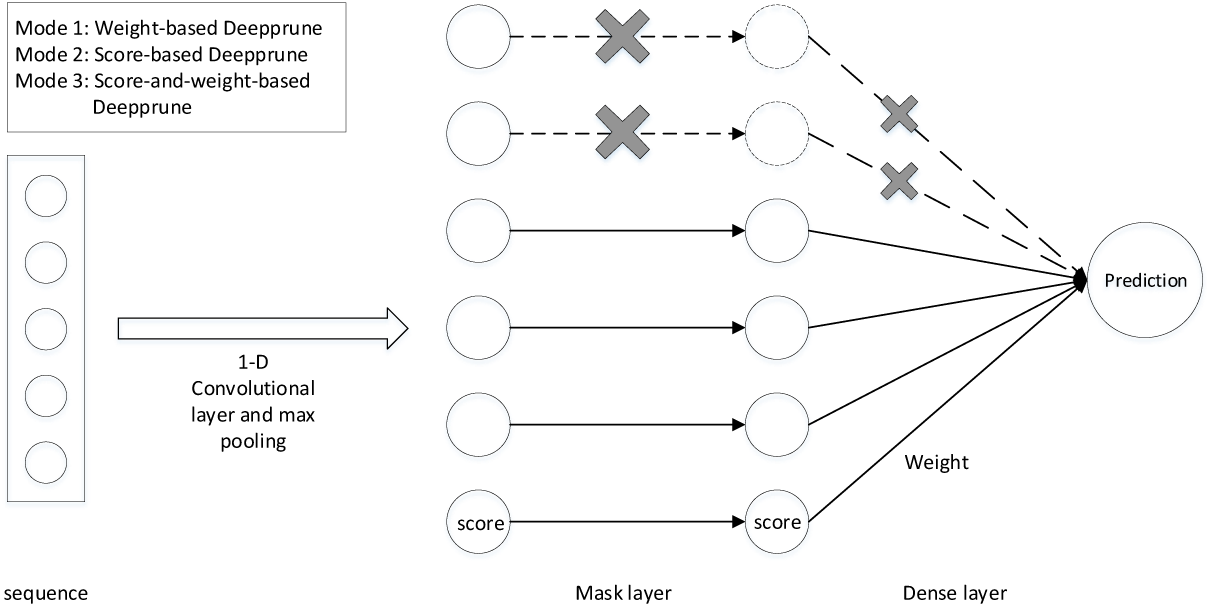
The architecture of Deepprune. The first layer is a convolutional layer. The second layer is a rectified linear units activation function followed by global max pooling. A mask layer is added to prune the small-magnitude weights. The fourth layer is a dense layer which linearly combines the outputs of all the kernels. The last layer is a sigmoid activation function which converts the values obtained in the dense layer to a value between 0 and 1 which corresponds to a probability.

**Figure 2.**
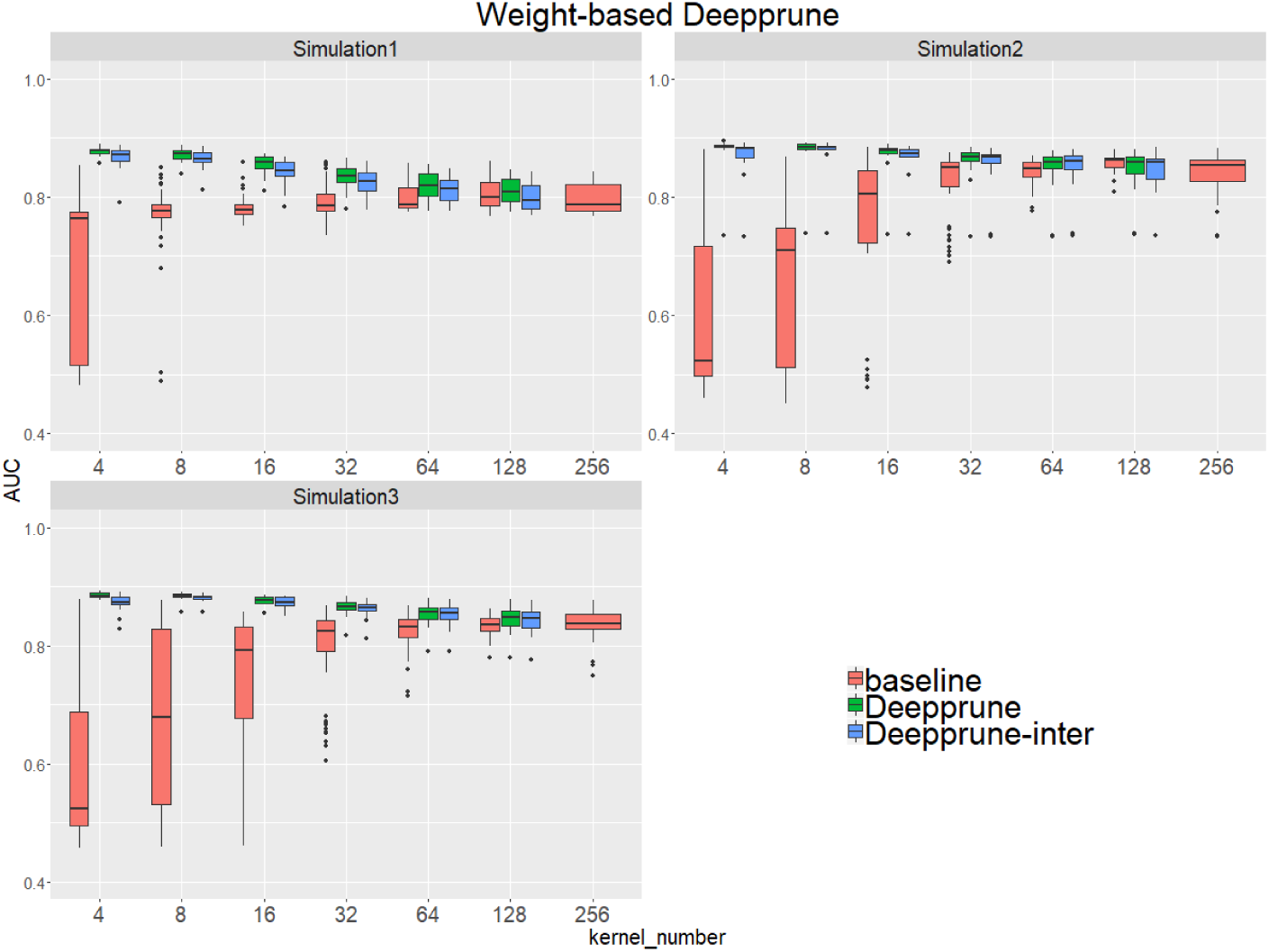
Weight-based Deepprune performs much better and is much robuster to different random initialization than baseline when kernel number is limited in the first three simulated datasets. The x axis shows the kernel numbers utilized in the model, and the y axis shows the AUCs obtained in testing. As kernel number decreases, the performance of Deepprune shows a converse tendency compared to baseline; thus, as iteration continues in an upward gradient, the mean AUC of Deepprune significantly improves.

Compared with the baseline, performance improvement was notably evident on the simulated dataset 4 and 5 with a hard true model, reflecting the excellence of Deepprune in cases with the complex motif settings [Figure 3]. Distinctly, the performance of baseline with 4 kernels is close to that of random guess on the complex datasets. This result shows that the baseline model with limited kernel numbers does not satisfy the need for overcoming the local optimum problem and that it lacks robustness to initialization. To our surprise, when the kernel number is half that of the motif number, the performance of Deepprune only drops a little, showing that the condition *d* = 4 is enough. What’s more, fewer kernels helps to improve the interpretation of our model. We also find that Deepprune-inter always shows poorer results, no matter whether from the mean AUC or the variation of AUC, which demonstrates that fine-tuning is essential in Deepprune.

**Figure 3.**
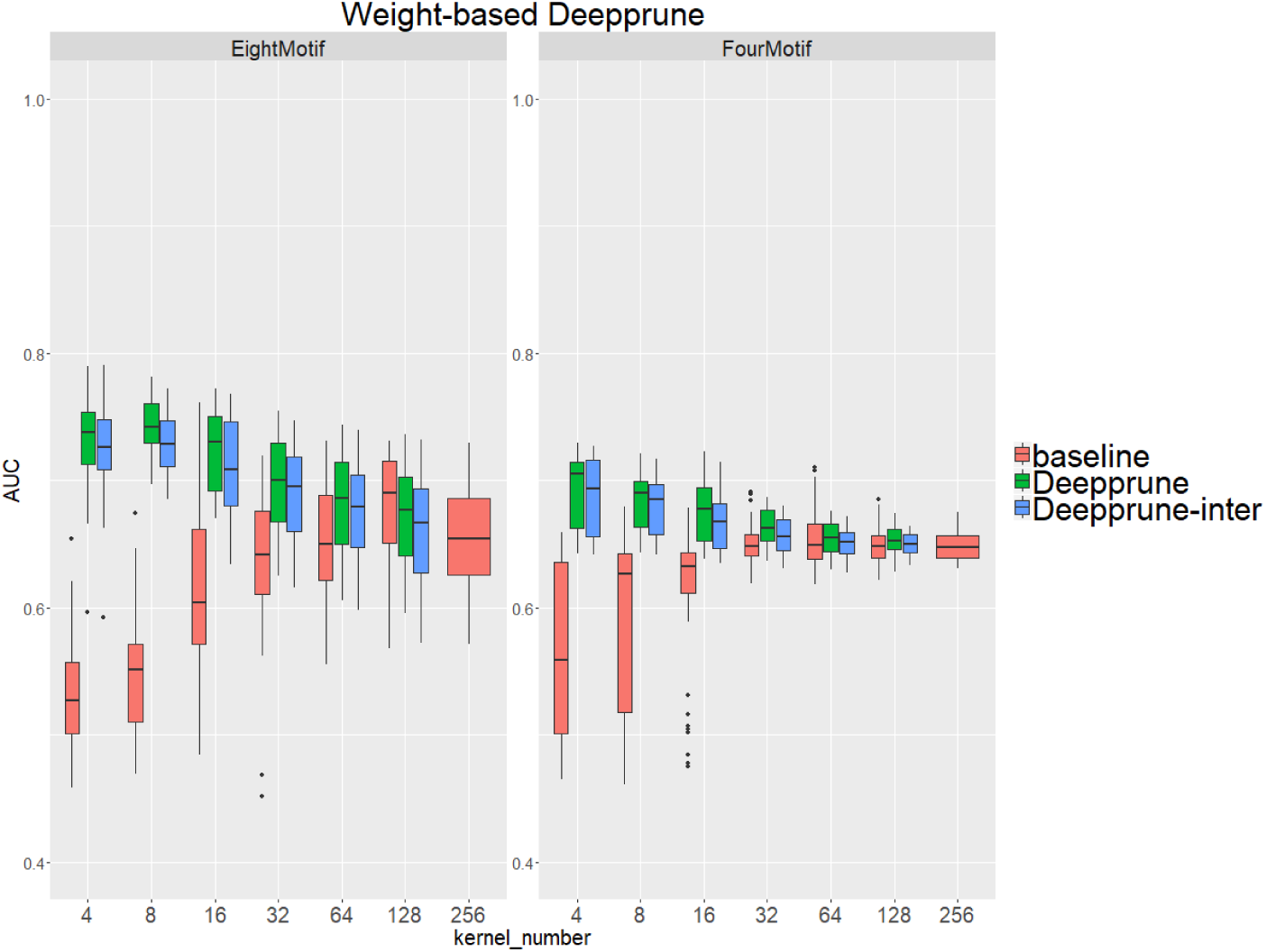
Weight-based Deepprune performs much better and is much robuster to different random initialization than baseline when kernel number is limited in the last two complex simulated datasets, even when kernel number is half the motif number at which time the performance of Deepprune only drops slightly.

### Comparison of three pruning criteria

Next, we studied the effects of the three criteria on the performance of Deepprune, as noted previously. We selected three simulated datasets to determine the difference of three different rules. If the scores are considered when pruning, then all samples in the training set need to be calculated, which leads to substantial calculation.

From Figure 4, when kernel number is high (e.g., 8 and 16), the performance of the three methods is nearly identical. Thus, the choice of there pruning methods is not crucial because the restriction to the kernel number is loose. However, when the kernel number is extremely limited, weight-based Deepprune shows its superiority compared to the other two methods in simulated dataset 1, in which the samples are hard to classify because of the information entropy in the true model. It is likely that weight-based Deepprune does not depend on samples which may cause randomness. From the case study below, the weights in the dense layer have a close magnitude, indicating that the scaling problem of scoring is difficult to solve in the smooth training process. Based on this observation, we select weight-based Deepprune as default.

**Figure 4.**
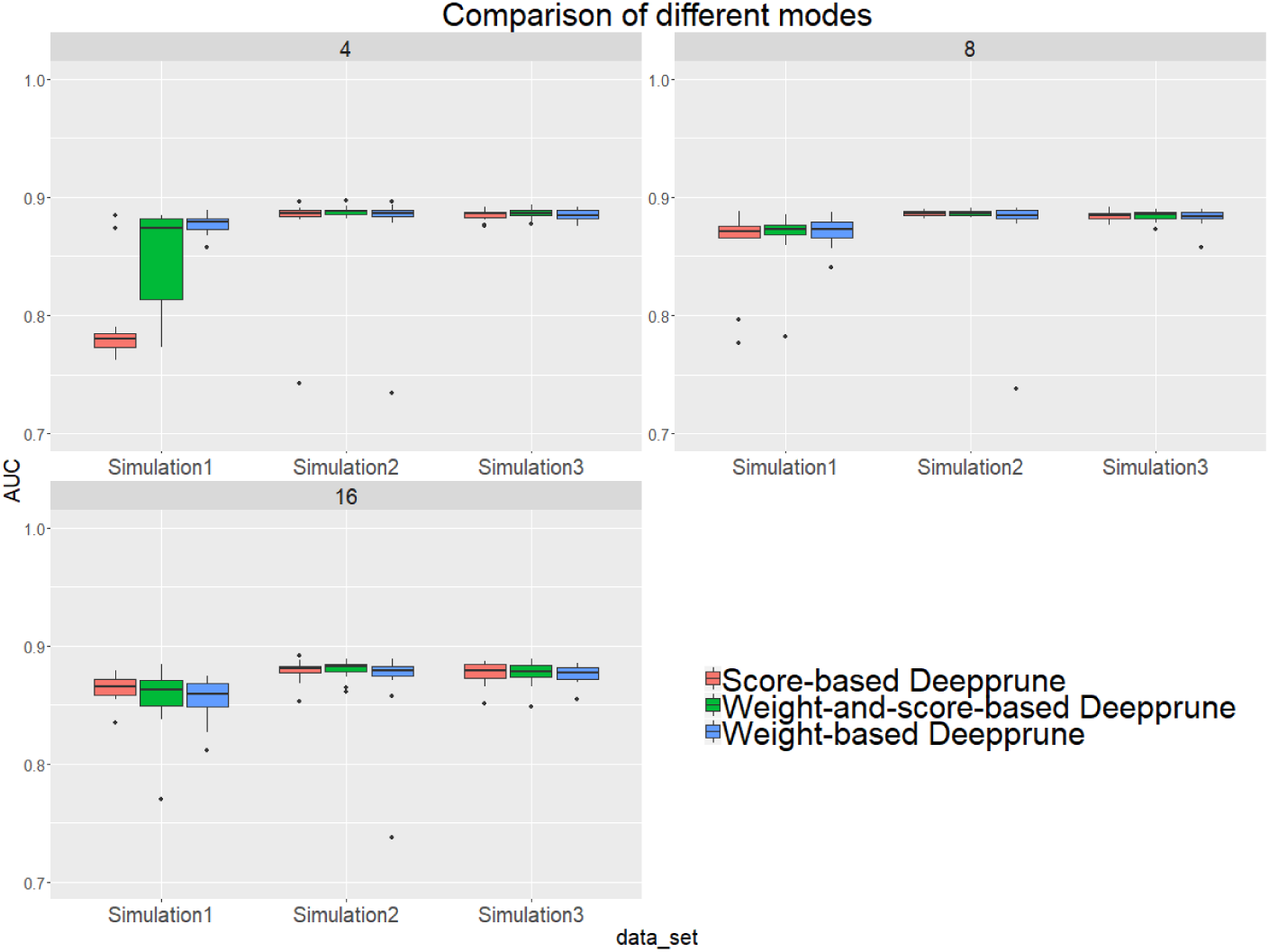
Three models of Deepprune are compared on the simulated datasets. We show the performance of Deepprune based on different criteria with 4, 8 and 16 kernels. The performance of three different criteria is almost identical to that with 8 and 16 kernels during iterative pruning. However, the final model with 4 kernels shows that weighted-based Deepprune is superior to the other two methods in simulated dataset 1, but hard to classify owing to high entropy.

### Performance on real datasets

We test the performance of DeepPrune on read data analysis in this section. CNN parameters are set the same as those for the simulated datasets, except the kernel length was changed to 15.

When the number of kernels is limited (i.e., 4), Deepprune achieves a statistically significant improvement in AUC from one-sided Wilcoxon signed-rank test in Figure 6, p=1.02×10*^−^*^58^, with a better performance on 77.10% of the datasets [Table 2]. Nevertheless, its accuracy is lower on 22.90% of the datasets, which does not match our expectation. This may be due to the non-convexity of the neural network model where local optimum is obtained. So we select the datasets for which our model’s performance is lower, and we initialize the training with several different random seeds.In some of the selected datasets, the mean performance of DeepPrune is almost as good as the baseline (see Supplementary Material). However, a consistent gap still appears in a small number of datasets in which the baseline shows better performance than our method, suggesting that the interaction of motifs is not considered in our architecture. It follows that the proposed architecture cannot represent the true model for some proteins in motif inference, which, therefore, creates bias for Deepprune.

**Table 2.** Performance of Deepprune on real data

| Kernel number | Method | AUC | Percentage improved | P-value |
| --- | --- | --- | --- | --- |
| 4 | baseline | 0.7785 |  |  |
|  | Deepprune | 0.8016 | 0.7710 | 1.02E-58 |
| 8 | baseline | 0.8169 |  |  |
|  | Deepprune | 0.8288 | 0.7174 | 6.44E-38 |
| 16 | baseline | 0.8432 |  |  |
|  | Deepprune | 0.8476 | 0.6826 | 2.63E-21 |
| 32 | baseline | 0.8602 |  |  |
|  | Deepprune | 0.8625 | 0.6681 | 3.25E-15 |
| 64 | baseline | 0.8728 |  |  |
|  | Deepprune | 0.8743 | 0.6507 | 1.20E-15 |
| 128 | baseline | 0.8809 |  |  |
|  | Deepprune | 0.8820 | 0.6986 | 4.41E-26 |
| 256 |  | 0.8849 |  |  |

### Case study

We selected several kernels to track the change of their corresponding weights at different pruning stages in the dense layer. In this section, we utilized simulated dataset 3 for we only knew the true models in simulated datasets. We chose the weights of 4 unpruned kernels and 2 pruned kernels at the end of each fine-tuning step. All the weights were collected after fine-tuning. It should be noted that the weights of the kernels in the convolutional layer changed during fine-tuning.

From Table 3, we can see that the magnitude of weights is gained step-by-step for four unpruned kernels, indicating that kernels show their importance over a gradual upward gradient. Before pruning, the weights of unpruned kernels are scrapped by auxiliary kernels. After pruning auxiliary kernels, the weights of unpruned kernels aren’t affected any more, which shows the superiority of Deepprune.

**Table 3.** Absolute value of weights of several kernels during different pruning stages in the dense layer.

| kernel number | kernel 1 | kernel 2 | kernel 3 | kernel 4 | kernel 5 | kernel 6 |
| --- | --- | --- | --- | --- | --- | --- |
| 256 | 0.9150 | 0.9019 | 0.7043 | 0.8112 | 0.2192 | 0.3582 |
| 128 | 0.9462 | 0.9294 | 0.7322 | 0.8344 | 0.2625 | 0.4015 |
| 64 | 0.9548 | 0.9364 | 0.7305 | 0.8252 | 0.2100 | 0.3750 |
| 32 | 0.9616 | 0.9403 | 0.7540 | 0.8387 | 0.0000 | 0.4433 |
| 16 | 0.9836 | 0.9504 | 0.8153 | 0.8589 | 0.0000 | 0.4584 |
| 8 | 1.0919 | 1.0501 | 0.9620 | 1.0249 | 0.0000 | 0.0000 |
| 4 | 1.2832 | 1.1979 | 1.2065 | 1.2542 | 0.0000 | 0.0000 |

### Model visualization

Now we study the ability of Deepprune to recover the underlying motifs more accurately. As in the last section, we utilized simulated dataset 3 because we only knew the true motifs in simulated datasets. The sequence logos are generated from kernels the way introduced in Section 10.2 of the DeepBind(Alipanahi et al., 2015) Supplementary Materials. The two best-recovered motifs, from the perspective of information content, were compared to the true motifs utilized on the simulated data. Their similarity (E-value) were also calculated utilizing the Tomtom algorithm(Gupta et al., 2007).

In Figure 5 the motifs recovered by Deepprune and the baseline were both aligned to the true motifs. We clearly found the sequence logos generated by Deepprune were informative and accurate from the E-value. The base-recovered motif by the baseline with 4 kernels exhibited very bad performance and the short motif in simulated dataset 3 could not be matched by 4 filters. In addition, we found that the motif regions could be distinguished from other regions which clearly obey background distribution. Although the length of kernels is far beyond that of the true motifs, the extra positions, which are not aligned to the true motifs, do not contain any noise, owing to the ability of Deepprune to lessen the impact of auxiliary kernels at the end of training.

**Figure 5.**
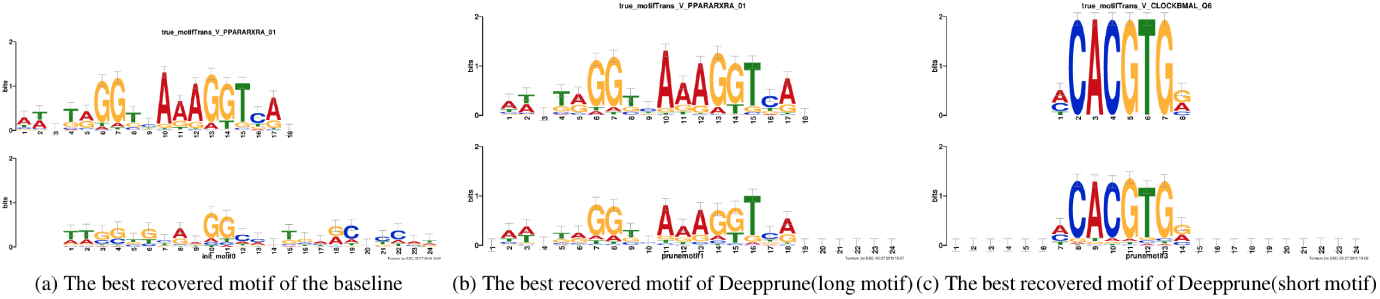
Motifs recovered by our model (the last two) and by baseline (the first) aligned to the true given motifs (top row). Utilizing Tomtom algorithm, the E-values of the motifs recovered by Deepprune are 1.35 *×* 10*^−^*^23^, 2.27 *×* 10*^−^*^7^, respectively. At the same time, the ones recovered by baseline are 1.64 *×* 10*^−^*^2^.

**Figure 6.**
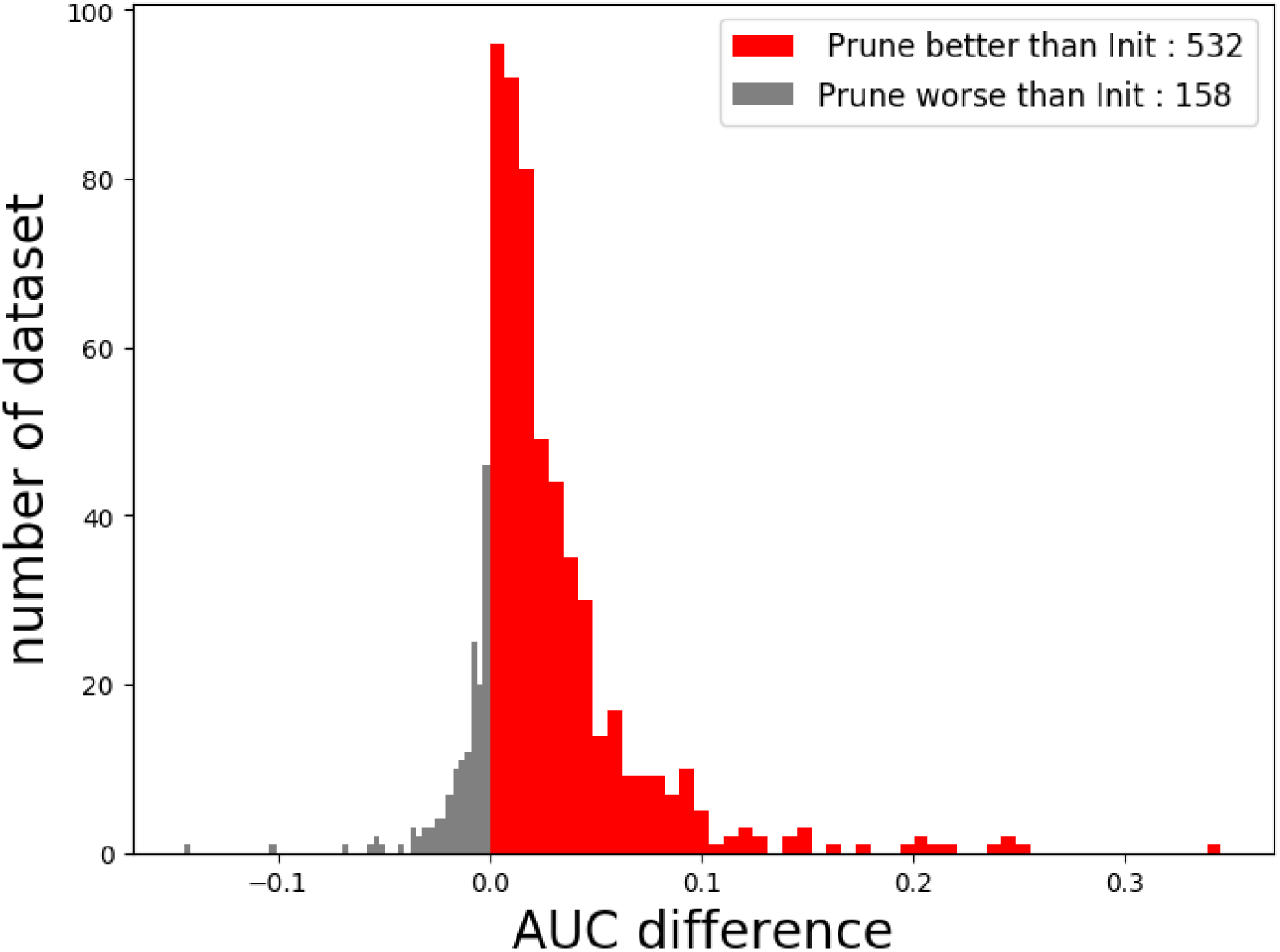
The performance of Deepprune with 4 kernels on real datasets where kernel length = 15. Deepprune greatly increases the AUC for real datasets. The AUC difference under the baseline (Init) and Deepprune (Prune) is shown from the x axis. Deepprune is better than baseline on 532 datasets, but worse than baseline with 158 datasets. This figure clearly shows that Deepprune achieves better performance with limited kernel number.

## DISCUSSION

### Regularization behind Deepprune

*L*_0_, *L*_1_ and *L*_2_ regularizations are three significant shrinkage methods for variable selection, and they are widely utilized in deep learning(Luo et al., 2019; He et al., 2016; Liu et al., 2017). However, the architecture of deep learning is multilayered and complex. Thus, for the same result, all weights in the architecture have the same infinite solution, e.g., the scaling problem noted before. *L*_1_ and *L*_2_ regularization update the original loss function by adding differentiable regularization terms, while *L*_0_ regularization needs to be realized by pruning. Actually, Deepprune adds *L*_0_ regularization to the weight in the dense layer instead of the entire architecture. Iterative pruning can help avoid wrong pruning for the greedy algorithm compared with one-shot pruning, thus showing its superiority in many tasks(Frankle and Carbin, 2018). Although *L*_1_and *L*_2_ penalties have been added to our model, the result shows little difference.

### Deep models are necessary for modeling TF-DNA specificities

Blum and Kollmann(Blum and Kollmann, 2019) supposed that deep models may be unnecessary for modeling TF-DNA specificities because they think that biological sequences are not composed of complex hierarchies of patterns as those in images. Deepprune can improve the performance of motif inference on real-world data compared with baseline, even with the same kernel number. However, since the weights are pruned iteratively, the performance of Deepprune does not change as what we saw in the simulated datasets. If PWM characterizes the specificities of motif inference and motif relationships are the same with those in simulated datasets, we will most likely see consistent performance in real-world and simulated datasets. In actuality, however, about 23% of datasets have a decrease compared to baseline with 4 kernels. As a result, we suspect that the interaction of different motifs and other complex relationships corresponding to motif inference need to be considered. Actually we suggest using different architectures to model different protein-binding problems. It is clear that adding the hidden layer gives deep learning architectures the ability to represent the interaction of different motifs and sequences of recurrent neural network models from the viewpoint of natural language processes, allowing various representations with different parameters. However, based on the results of our experiment, many biological sequences cannot be modeled very well by the simple DeepBind model, making it necessary to create deeper architectures to identify the underlying model for some proteins.

### Lottery Ticket Hypothesis

Recently, the lottery ticket hypothesis has attracted attention in the field of deep learning. This hypothesis holds that dense, randomly initialized networks contain subnetworks that, when trained in isolation (i.e., utilizing the same initialization), reach test accuracy comparable to the original network in a similar number of iterations(Frankle and Carbin, 2018). The subnetworks are called winning tickets. Liu et al.(Liu et al., 2018) even suggested that the value of automatically structured pruning algorithms sometimes lies in identifying efficient structures and performing implicit architecture search, rather than selecting “important” weights, irrespective of the initiation. First, if the architecture of our pruned model is equal to that of baseline, our result in the simulated dataset shows that either weight or initiation also counts for the performance of unstructured pruning algorithms. Second, we tried to find our winning tickets by following the methods in the original paper. We substituted the fine-tuning step in Deepprune for retraining, which resets the remaining parameters to their values before initial training. Experiments on real data with winning tickets realize slightly better performance (Mean AUC is 0.8035 with 4 kernels), which shows that this hypothesis may be true for Deepprune (see Supplemental Material).

## CONCLUSION

In this study, we proposed a novel deep-learning framework called Deepprune, to improve the performance of predicting the binding preference of DNA-protein binding. Deepprune prunes weights of kernels in the dense layer and fine-tunes iteratively by adding a mask layer in the architecture of motif inference. Deepprune utilizes limited kernel number in the convolutional layer, which shows the efficiency and interpretability of our model. In this study, Deepprune is shown to improve model performance compared with baseline with the same limited kernel number, both in simulated and real-world datasets. Our method improves performance without changing the basic architectures or adding extra parameters at the end of training.

To the best of our knowledge, we are the first to introduce a pruning framework in the field of motif inference. Network pruning has been widely applied in the framework of deep learning for its ability to reduce storage and computation without affecting accuracy. Although the architecture of motif inference is very simple, network pruning is meaningful for the model since the use of fewer kernels can still achieve better interpretation as can be seen from model visualization.

The motif-finding problem remains unsolved. Deep learning is very useful for complex structures and large datasets. What’s more, it has greatly improved the state-of-the-art in many areas. Neural networks have achieved a lot of success, such as DeepBind and DeepSEA for motif-finding. However, in spite of the great achievements, deep learning is blamed due to the lack of interpretability as well(Castelvecchi, 2016; Zou et al., 2018). DeepBind shows its superiority compared with conventional machine learning methods, which proves that both deep and complex representation of the sequences is essential for motif inference. Because the gap between the performance on simulated and real-world datasets, we wonder if this is due to the underlying model behind some of the real-world datasets is complex.

Recently, many studies have investigated the interpretation of neural networks and the underlying model behind real-world datasets. They utilize complex models, such as RNN and the model with attention mechanism, which comes from the field of natural language processing, to represent the complex information of biological sequences(Zuallaert et al., 2018; Luo et al., 2019; Shen et al., 2018; Pan and Shen, 2018; Pan and Yan, 2017; Li et al., 2019; Pan et al., 2018). Actually, from the diversity of DNA-protein binding, we suggest using different architectures to model motif inference for specific proteins. Complex network architectures combined with pruning technology can result in approximating the true model of motif inference. Since our basic architecture is simple, adding a hidden layer before the dense layer and then adding an RNN layer after the convolutional layer, as well as replacing global max pooling with expectation pooling, can also be considered, but these topics are outside the scope of the present paper.

## AUTHOR CONTRIBUTIONS

XL, WC and MD designed the experiments; XL drafted the manuscript; WC carried out the experiments; XL and WC analyzed the results. All authors read and approved the final manuscript.

## ACKNOWLEDGMENTS

None

## SUPPLEMENTAL DATA

See Supplemental Material.

## DATA AVAILABILITY STATEMENT

All code is public and can be found at https://github.com/klovbe/Deepprune

