## Supplemental Material for "Deepprune: Learning efficient and interpretable convolutional networks through weight pruning for predicting DNA-protein binding"

### *Supplementary Material*

#### 1 SUPPLEMENTARY FIGURES

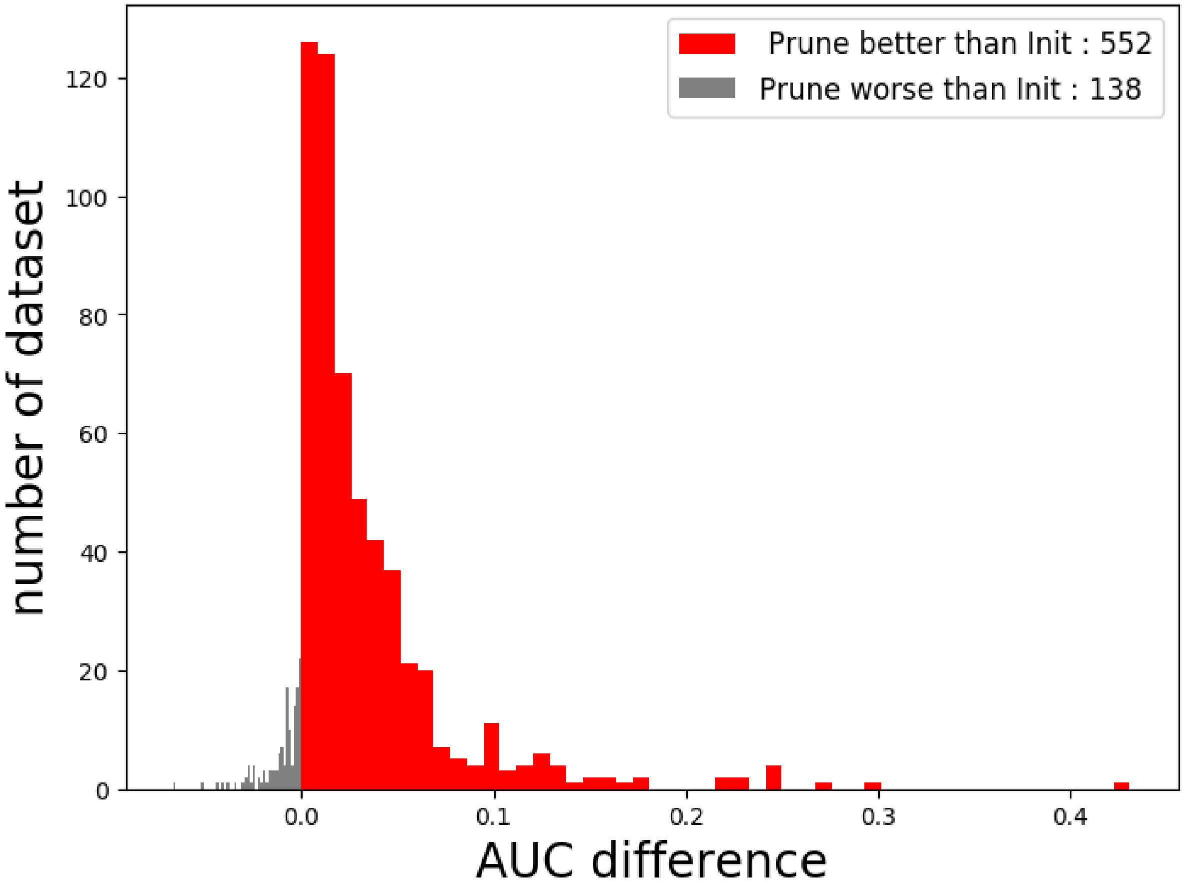


**Figure S1.** The performance of Deepprune with reinitialization of winning tickets on real datasets where kernel length = 15 and kernel number = 4. Deepprune with reinitialization of winning tickets increases the AUC on real datasets. The x axis shows the AUC difference under the baseline (Init) and Deepprune (Prune). Deepprune with re-initialization is better than baseline on 552 datasets, but worse than baseline with 138 datasets. This figure clearly shows that Deepprune with reinitialization of winning tickets achieves better performance with limited kernel number.

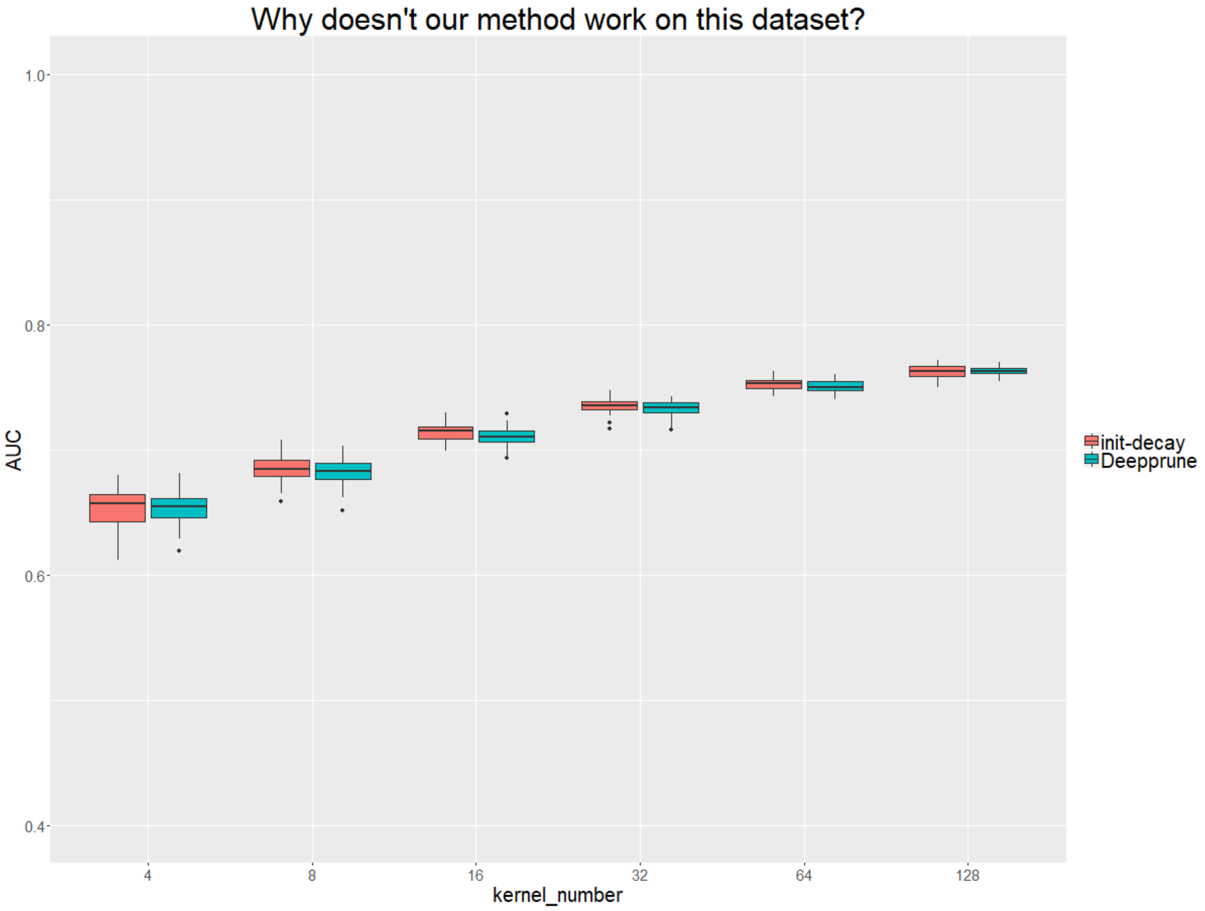


**Figure S2.** We selected the datasets in which the performance of Deepprune was below 0.01 and executed them 10 times with different random seeds to determine whether the poor performance was consistent. As shown, on these datasets, Deepprune is not always worse than the baseline; in fact, the mean performance of the two models is almost identical. However, we also found that our model performances worse in part of the datasets. We suspect that the underlying model of these datasets is complex.
